# Analysis of the genome-scale metabolic model of *Bacillus subtilis* to design novel in-silico strategies for native and recombinant L-asparaginase overproduction

**DOI:** 10.1101/2022.12.29.522229

**Authors:** Nisha Sanjay Barge, Ansuman Sahoo, Veeranki Venkata Dasu

## Abstract

L-asparaginase is an enzyme with widescale use in the food and medicine industry. It is used as a chemotherapeutic drug in the treatment of acute lymphoblastic leukemia. Limitations of side effects associated with commercially available L-asparaginase necessitate the search for alternative sources. *Bacillus subtilis* is an emerging host for the production of chemicals and therapeutic products. This study deals with L-asparaginase production in *Bacillus subtilis* using systems metabolic engineering approach. System biology offers a detailed understanding of organism metabolism at the network level unlike the conventional molecular approach of metabolic engineering allowing one to study the effects of metabolite production on growth. Metabolism of *Bacillus subtilis* is studied using genome-scale metabolic model iYO844 which consists of relationships between the genes and proteins present in *Bacillus subtilis*. Also, the model contains information about all the metabolic reactions and pathways allowing convenient metabolic engineering methods. Computational methods like flux balance analysis, flux variability analysis, robustness analysis, etc. are carried out to study the metabolic capabilities of *Bacillus subtilis*. The model predicted a specific growth rate of 0.6242 h^-1^, which was comparable to the experimental value. Further, the model is used to simulate recombinant L-asparaginase production generating a maximum production rate of 0.4028 mmol gDW^-1^ h^-1^. Flux scanning based on enforced objective flux and OptKnock design strategies are used for strain development of *Bacillus subtilis* for higher production of both native and recombinant L-asparaginase.

## 1. Introduction

Acute lymphoblastic leukemia (ALL), a type of blood cancer, affects the lymphoid cell line leading to the abnormal proliferation of lymphoid progenitor cells and the development of immature lymphocytes. ALL majorly affects children and young adults and is considered as the most common pediatric cancer responsible for the highest mortality rate in cancers in children [1]. Although considered as a highly curable cancer in the case of pediatric patients, the overall cure rate drops down to 40% in adults because of higher drug resistance, lesser tolerance to treatment, and less effective treatment regimen in adults compared to the pediatric population [2]. Amongst a range of therapies available comprising radiation therapy and immunotherapy; long-term chemotherapy is currently the most widely used treatment for ALL. The chemotherapeutic drugs involved in the therapy include prednisone, dexamethasone, methotrexate, asparaginase and mercaptopurine (during consolidation), vincristine, cytarabine, doxorubicin, etc. Amongst these, pediatric patients administered with asparaginase enzyme possess a higher survival rate compared to the patients not treated with asparaginase [3]. Since the first discovery of asparaginase as an anti-lymphocytic agent derived from guinea pig serum in 1953, asparaginase has been a major therapeutic drug in the treatment of ALL and is prominently produced from two bacterial sources - *Escherichia coli* and *Erwinia chrysanthemi* [4]. Although highly effective, asparaginase administration during ALL treatment derived from *Escherichia coli* is associated with several side effects including allergic reactions, development of antibodies, liver dysfunction, hyperglycemia, etc. [5,6]. *Erwinia* asparaginase and PEGylated form of asparaginase are considered as suitable substitutes which mitigate the risk of hypersensitivity to some extent though not entirely [7,8]. The side effects of the highly efficient asparaginase treatment necessitate the search for alternative asparaginase enzyme sources with lesser or completely devoid of side effects.

*Bacillus subtilis* has emerged as a potential cell factory platform for the production of recombinant proteins relevant to the pharmaceutical industry [9]. Recently, the anticancer activity of L-asparaginase isolated from a *Bacillus subtilis* species was tested and found to be effective against liver, breast, and colon cancer cells. Further, the isolated L-asparaginase showed no glutaminase activity which is responsible for the side effects of previously described asparaginases [10]. This article discusses the study dealing with the production of L-asparaginase from *Bacillus subtilis* 168 species using systems metabolic engineering approach. Ever since the rapid advancements in genome sequencing and bioinformatics tools, a large amount of high throughput omics data is made available which can be used for several biotechnological applications. Genome-scale metabolic models (GSMM) have emerged as an important tool for understanding the metabolism of an organism at a systems level. GSMM are employed in metabolic engineering and strain improvement targeted at overexpression of a particular metabolite [11]. GSMM allows the use of various analysis methods like flux balance analysis, flux variability analysis, gene essentiality analysis, in silico gene knockout targets, etc. which find wide applications in metabolite production from microbes, drug targeting in pathogens, enzyme function predictions, studying interactions between multiple cells and organisms and so forth [12]. The study carried out in this report represents the attempts to overexpress native L-asparaginase in *Bacillus subtilis* 168 using the already reconstructed and experimentally validated genome-scale metabolic model iYO844. Initially, the model is analyzed to study *Bacillus subtilis* metabolism under various growth conditions. The analysis is then used to design overexpression strategies for the production of native L-asparaginase by identifying gene amplification and knockout targets. Further, the model is analyzed to overexpress recombinant L-asparaginase gene from *Rhizomucor miehei* to produce glutaminase-free L-asparaginase and design strategies are designed for the same.

## 2. Methodology

### 2.1 Genome-scale metabolic model analysis

The genome-scale metabolic model of *Bacillus subtilis* titled iYO844 was downloaded in SBML format from the biochemical network reconstruction database maintained by Systems Biology Research Group (https://systemsbiology.ucsd.edu/InSilicoOrganisms/Bsubtilis/BsubtilisSBML). The model is then integrated into MATLAB version R2019b using COBRA Toolbox version 2021. The functions employed to carry out the further discussed objectives using the MATLAB platform are mentioned in detail in Supplementary file S1. Biomass growth is the objective function for iYO844. The model growth is simulated in glucose minimal growth media with glucose as a carbon source. The rest of the constituents include Fe^3+^, calcium, sulfate, potassium, magnesium, ammonia, and phosphate. The growth is simulated in aerobic conditions hence oxygen flux is unconstrained. To replicate experimental conditions, the bounds on substrate and oxygen exchange reactions are set as per requirement. The bounds of the required reaction are changed using the ‘*changeRxnBounds’* function in COBRA Toolbox. The specific glucose uptake rate for *Bacillus subtilis* 168 in glucose minimal media was set to 15 mmol gDW^-1^ h^-1^ [13] and the specific oxygen uptake rate was set to 20 mmol gDW^-1^ h^-1^ according to the value experimentally obtained for *E. coli* fermentation [14]. The model is then primed for further analysis including flux balance analysis, flux variability analysis, and robustness analysis.

### 2.2 Flux balance analysis (FBA)

FBA calculates the flow of metabolites through this metabolic network assuming steady-state conditions, thereby making it possible to predict the growth rate of an organism or the rate of production of a biotechnologically important metabolite [15]. The biomass growth predicted corresponds to the specific growth rate in the exponential phase denoted by μ with units per time. FBA is performed by using the in-built COBRA Toolbox ‘*optimizeCbModel* function. The objective function can be changed from biomass growth function to any other reaction of interest present in the model to predict metabolite production rate. Performing FBA generates the result including the value of the objective function, the time elapsed to perform the analysis, and a vector comprising the flux values of all the reactions present in the model. Usage of the function is discussed in Supplementary file S1.

### 2.3 Flux variability analysis (FVA)

The constraint-based optimization analysis used for analyzing the GSMM generates multiple alternate optimal solutions. The alternate optimal solutions represent situations where the system makes use of different sets of flux distributions to achieve the same objective function value. Due to the multiple solutions obtained for achieving the same biomass growth rate through different flux distributions, it becomes necessary to know the range of flux values a particular reaction can attain under specified conditions which are carried out by using the flux variability analysis approach. Theoretically, this is achieved by fixing the objective value and maximizing each reaction in the model followed by minimization to determine the feasible range of fluxes for every reaction [16]. FVA can be done for one particular reaction of interest or all the reactions present in the model. In practice, FVA is performed using the in-built function provided by COBRA Toolbox, *‘fluxVariability’* (detailed in Supplementary file S1). It is possible to determine essential, non-essential, flexible, and non-flexible reactions by analyzing FVA data. Reactions with non-zero lower flux limits are considered to be essential. Non-essential reactions have zero as their lower flux limits. Flexible reactions have unequal lower and upper limits and hence can carry a range of fluxes to meet the objective function while non-flexible reactions possess the same upper and lower limits and require a specific flux value to meet the objective function.

### 2.4 Robustness analysis

Robustness concerning metabolic networks is a measure of the change in the maximal flux of the objective function (here, biomass growth is considered as objective) when the optimal flux through any particular metabolic reaction is changed. Robustness analysis describes how biomass growth changes when the flux is increased through a particular reaction of interest [17]. Robustness analysis allows one to assess the ability of the host cell to adapt to the changes occurring due to metabolism (increase in flux through product reaction) or environmental changes. The function ‘*robustnessAnalysis’* generates a plot illustrating the change of objective flux along with the control reaction flux (specified by ‘controlRxn’). Usage detailed in Supplementary file S1.

### 2.5 Identification of gene amplification targets using flux scanning based on enforced objective flux (FSEOF) method

Flux scanning based on enforced objective flux (FSEOF) is a strategy used to identify amplification targets for desired metabolite production used initially for designing strain for lycopene production [18]. When flux towards product formation is added as an additional constraint on the model, FSEOF predicts a list of all the metabolic fluxes which increased along with product formation by scanning fluxes of all reactions present. The flux through native L-asparaginase (ASNN) reaction is gradually increased from 0 to 37 mmol gDW^-1^ h^-1^ with an interval of 5, as these are the lower and upper flux limits as obtained by FVA. After every step of setting a constraint on ASNN reaction to fix a particular flux value, FBA is carried out to obtain flux distribution (detailed in Supplementary file S1). The flux distribution for all ASNN values is scanned to determine the reactions which are gradually increasing and considered as reaction amplification targets. The interconnection of the target reactions with that of ASNN is visualized using in-built visualization functions.

### 2.6 Identification of gene knockout targets using OptKnock

OptKnock a computational method has been considered as a popular reaction deletion simulation algorithm used with GSMM. The algorithm is designed in such a way that it can predict a set of deletion targets that maximizes cell growth and the production of the target compound of interest using a bi-level optimization algorithm. OptKnock allows the selection of a number of reactions to be knocked out with default settings set at 5. The method was followed similar to the one used for the overproduction of biochemical products in *E. coli* [19]. The method is carried out by using the in-built *‘OptKnock* function in the COBRA Toolbox package. The input for the function includes specifying a target reaction to maximize. OptKnock also allows fixing a biomass growth value and predicts knockout targets at that particular biomass growth value. The result generates a list of reactions that can be knocked out for optimal growth and metabolite production rate.

### 2.7 Integration of heterologous L-asparaginase production metabolic reaction into iYO844 and design strategies for overproduction (FSEOF and OptKnock)

L-asparaginase produced by the fungal species *Rhizomucor miehei* was reported to accelerate the apoptosis of K562 leukemia cells indicating its anti-cancer activity. Further, a negligible amount of glutaminase activity was detected [20]. Thus L-asparaginase from *Rhizomucor miehei* was considered as a potential candidate in the quest for alternative drug sources for ALL. The gene for L-asparaginase was added to *Bacillus subtilis* iYO844 in the form of a reaction based on the amino acid sequence of *Rhizomucor miehei* L-asparaginase as detailed by the procedure in [21]. The reaction was designated as ‘rASNN’ representing recombinant L-asparaginase. Exchange reaction for rASNN was added to the model to simulate the extracellular expression of L-asparaginase production. The reactions were added using ‘*addReaction*’ in-built function available in COBRA Toolbox. The details of the construction of the metabolic reaction representing heterologous L-asparaginase enzyme production and the functions used are discussed in Supplementary file S1. The model is then analyzed by FBA, FVA, and robustness analysis after the addition of heterologous L-asparaginase production reaction. Over-production strategies for the same are designed using FSEOF and OptKnock methods as detailed in previous sections. Further details are mentioned in Supplementary file S1.

### 2.8 Adding amino acids to the media for increasing rASNN production

The production of rASNN was analyzed when an external source of amino acids is provided in the media. The conditions are simulated by adding exchange reactions for all 20 amino acids and are constrained by specifying the uptake rates specified for *E. coli* as mentioned in [22]. The uptake rates are set by altering the lower bound values to that of the values obtained in the literature (MATLAB commands mentioned in Supplementary file S1).

### 2.9 Visualization of metabolic networks

The metabolic networks can be visualized using the in-built functions in the COBRA Toolbox package, *‘draw_by_rxn*’. Initially, a cell vector containing the reaction IDs of reactions to be included in the layout is specified. The function generates a biograph figure illustrating the interconnections of the reactions specified. It also generates a list of metabolites present in the reaction pathways and dead-end metabolites. Implementation of the function is discussed in Supplementary file S1.

## 3. RESULTS AND DISCUSSION

### 3.1 Analysis of *Bacillus subtilis* iYO844 model

The genome-scale metabolic model contained 844 genes, 990 metabolites, and 1250 reactions. On integration with the MATLAB platform, the model is displayed as a single MATLAB structure with numerous fields detailing the properties of the model. The model consists of information regarding the number of genes, metabolites, and reactions along with their complete scientific names and IDs. The model is initially unconstrained, indicated by the relatively higher upper and lower bound values for the reactions contained in the double vectors represented by ‘ub’ and ‘lb’ respectively. Biomass growth is set as the objective function for flux balance analysis of the model.

#### 3.1.1 *In-silico* growth of *Bacillus subtilis* using different carbon sources

The iYO844 model is validated by analyzing the *in-silico* growth of the model using various carbon sources and comparing it with experimental results for *Bacillus subtilis* growth obtained through published articles. The model was analyzed for growth on acetate, arabinose, fructose, mannitol, fumarate, citrate, galactose, glucosamine, malate, mannose, succinate, sucrose, and xylose. The results are tabulated in Table 1.

**Table 1:**
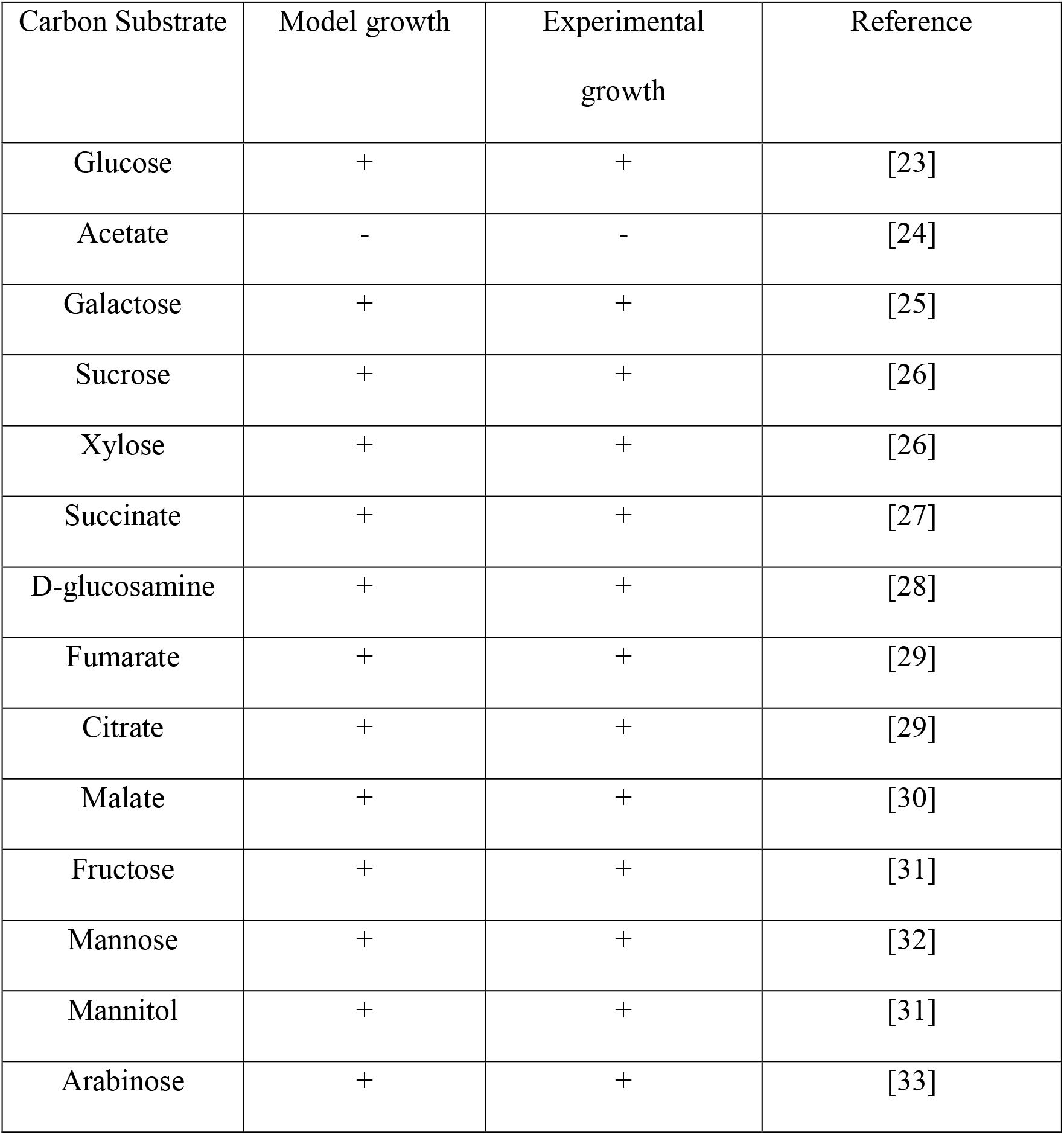
Validation of model growth on various carbon substrates in comparison with data obtained from the literature. (‘+’: indicates growth, ‘-’: indicates no growth)

| Carbon Substrate | Model growth | Experimental growth | Reference |
| --- | --- | --- | --- |
| Glucose | + | + | [23] |
| Acetate | - | - | [24] |
| Galactose | + | + | [25] |
| Sucrose | + | + | [26] |
| Xylose | + | + | [26] |
| Succinate | + | + | [27] |
| D-glucosamine | + | + | [28] |
| Fumarate | + | + | [29] |
| Citrate | + | + | [29] |
| Malate | + | + | [30] |
| Fructose | + | + | [31] |
| Mannose | + | + | [32] |
| Mannitol | + | + | [31] |
| Arabinose | + | + | [33] |

#### 3.1.2 Flux balance analysis (FBA) for iYO844

The specific growth rate of *Bacillus subtilis* for unconstrainted glucose and oxygen-specific uptake rates was found to be 0.118 hr^-1^. The flux balance analysis resulted in a glucose uptake rate of 1.7 mmol gDW^-1^ h^-1^ and an oxygen uptake rate of 5.706 mmol gDW^-1^ h^-1^. After setting the specific glucose uptake rate at 15 mmol gDW^-1^ h^-1^ and specific oxygen uptake rate at 20 mmol gDW^-1^ h^-1^ as detailed in section 2.1, the specific growth rate of *Bacillus subtilis* was found to be 0.6242 hr^-1^ which is comparable to the experimental growth rate obtained for glucose as the carbon source [23]. Flux through L-asparaginase (ASNN) reaction was zero. This is in correlation with the nature of microbial growth where product formation is minimized to favor bacterial growth.

#### 3.1.3 Flux variability analysis (FVA) for iYO844

FVA calculates the minimum and maximum allowed fluxes through all the reactions present in the metabolic network. Through FVA, it is found that the minimum flux through native L-asparaginase reaction is zero and the maximum allowed flux is 37 mmol gDW^-1^ h^-1^ under the given media conditions of glucose and oxygen constraints.

#### 3.1.4 Robustness analysis for native L-asparaginase (ASNN) reaction

Robustness analysis allows us to study the change in microbial growth rate as the flux through ASNN increases. This helps in selecting a trade-off value to obtain an optimal ASNN flux value and an optimal biomass growth rate. As the flux through ASNN is increased, the specific growth rate steadily decreases as seen during the production of industrially relevant products (Fig.1). At the highest growth rate of 0.6242 h^-1^, ASNN flux is zero and reaches a maximum of 37 mmol gDW^-1^ h^-1^ at 0 growth.

**Fig.1.**
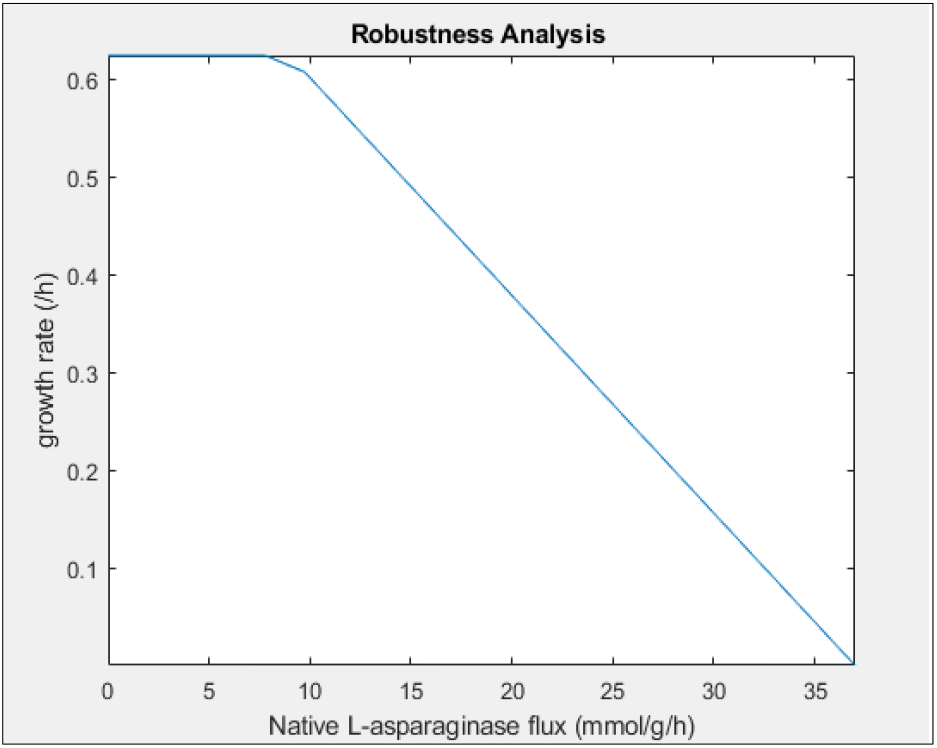
Plot representing the change of specific growth rate of *Bacillus subtilis* (h^-1^) with increase in native L-asparaginase reaction flux (mmol g^-1^ h^-1^) as obtained from robustness analysis

### 3.2 Strain design for native L-asparaginase (ASNN) overproduction

#### 3.2.1 FSEOF to identify reaction amplification targets for ASNN production

FSEOF identifies reactions for which the flux amplifies when flux through ASNN is gradually increased. The FSEOF method resulted in an increase in flux through the following listed reactions as flux through ASNN reaction was gradually increased from 0 to 35 mmol gDW^-1^ h^-1^ with an interval of 5 (highest value of 37 mmol gDW^-1^ h^-1^ is avoided as growth rate reduces to zero). In accordance with the results obtained from robustness analysis, the specific growth rate of *Bacillus subtilis* is seen to decrease as flux through the ASNN reaction is gradually increased (Fig.3). Of the total 21 potential reaction amplification targets, the reactions catalyzed by the respective enzymes tabulated in Table 2 are found to be metabolically relevant to the ASNN reaction. Fig.2 represents a biograph generated using MATLAB, illustrating the metabolic pathway connections of these reactions with that of ASNN.

**Fig.2.**
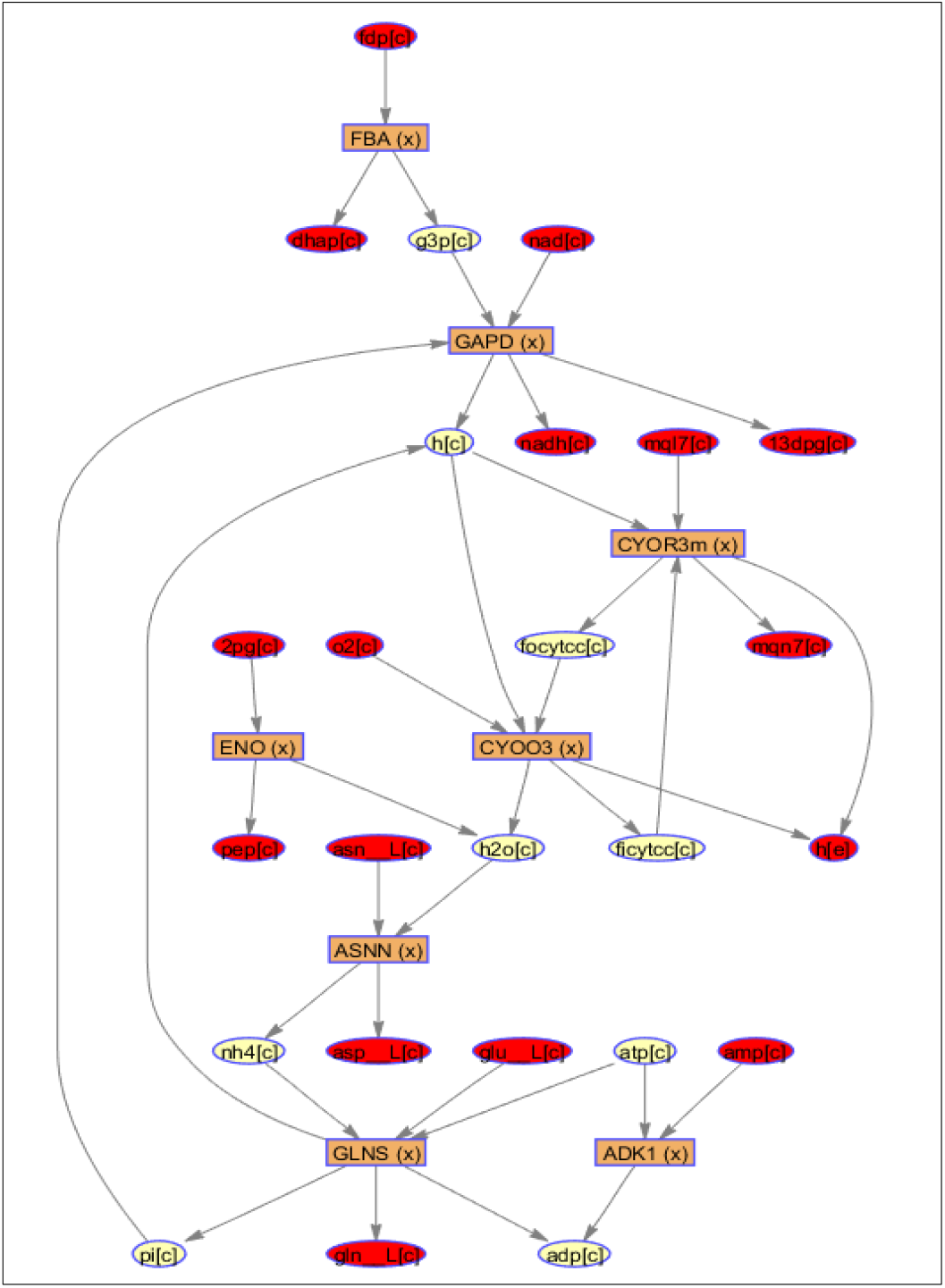
Biograph of metabolic network representing pathway connections of ASNN reaction with the identified amplification targets as generated by MATLAB R2019b in hierarchical layout

**Fig.3.**
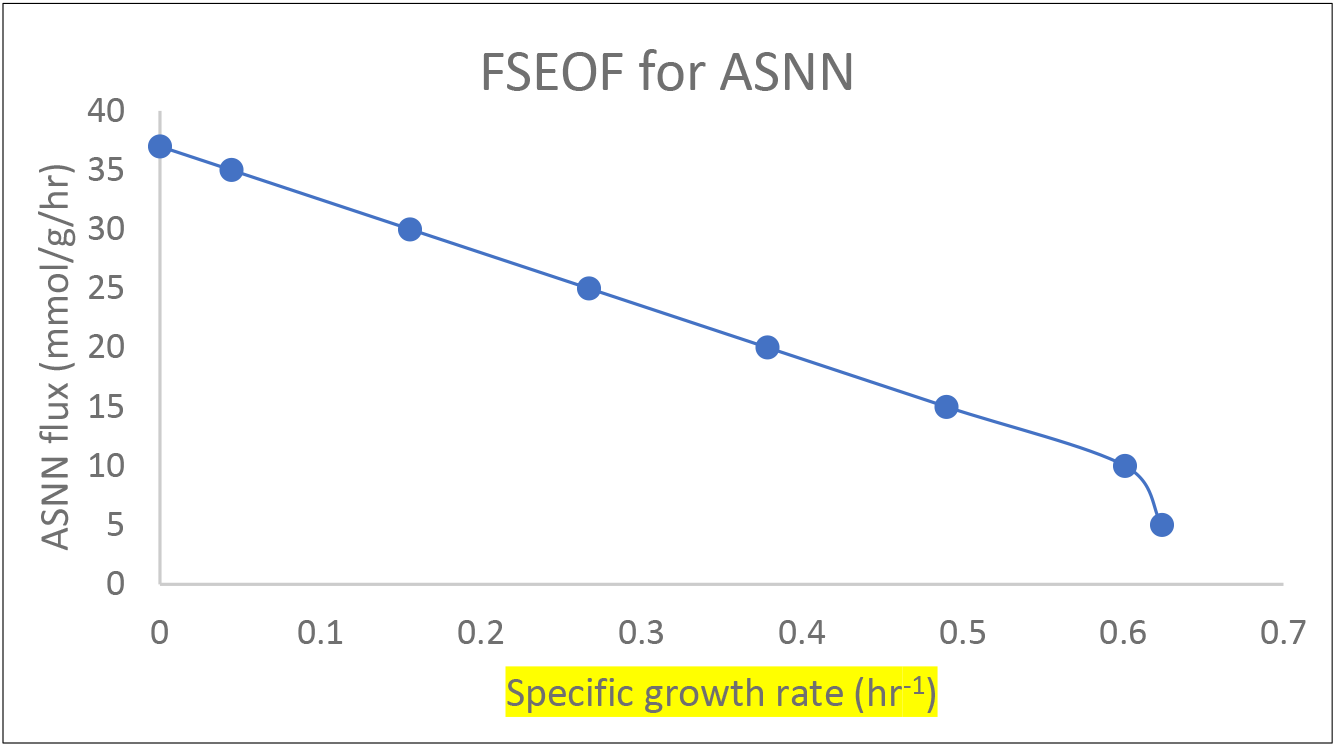
Plot representing the change in specific growth rate of *Bacillus subtilis* (h^-1^) as flux through ASNN reaction (mmol g^-1^ h^-1^) is gradually increased from the minimum to maximum limit as predicted by FVA. Growth rate increases linearly until 0.6 h^-1^ as ASNN flux decreases and reaches a maximum at 0 ASNN flux

**Table 2:** Reaction amplification targets identified by FSEOF for maximizing flux through native L-asparaginase reaction. The table contains reaction ID as present in the model, complete reaction name, and reaction formulae

| Reaction ID | Reaction name | Reaction formula |
| --- | --- | --- |
| FBA' | 'Fructose-bisphosphate aldolase' | $\text{fdp}[\text{c}] \rightleftharpoons \text{dhap}[\text{c}] + \text{g3p}[\text{c}]$ |
| ADK1' | 'Adenylate kinase' | $\text{amp}[\text{c}] + \text{atp}[\text{c}] \rightleftharpoons 2 \text{ adp}[\text{c}]$ |
| CYOR3m' | 'Cytochrome c reductase menaquinol 7 3 protons ' | $2 \text{ ficytcc}[\text{c}] + \text{h}[\text{c}] + \text{mql7}[\text{c}] \rightarrow 2 \text{ focytc}[\text{c}] + 3 \text{ h}[\text{e}] + \text{mqn7}[\text{c}]$ |
| CYOO3' | 'Cytochrome c oxidase He 2 ' | $2 \text{ focytc}[\text{c}] + 6 \text{ h}[\text{c}] + 0.5 \text{ o2}[\text{c}] \rightarrow 2 \text{ ficytcc}[\text{c}] + \text{h2o}[\text{c}] + 4 \text{ h}[\text{e}]$ |
| GAPD' | 'Glyceraldehyde-3-phosphate dehydrogenase' | $\text{g3p}[\text{c}] + \text{pi}[\text{c}] + \text{nad}[\text{c}] \rightarrow \text{13dpg}[\text{c}] + \text{h}[\text{c}] + \text{nadh}[\text{c}]$ |
| GLNS' | 'Glutamine synthetase' | $\text{atp}[\text{c}] + \text{glu\_L}[\text{c}] + \text{nh4}[\text{c}] \rightarrow \text{adp}[\text{c}] + \text{gln\_L}[\text{c}] + \text{h}[\text{c}] + \text{pi}[\text{c}]$ |
| ENO' | 'Enolase' | $2\text{pg}[\text{c}] \rightleftharpoons \text{h2o}[\text{c}] + \text{pep}[\text{c}]$ |

The results obtained from both FSEOF and OptKnock methods are analyzed based on the hypothesis that the rate of reaction (flux) increases with an increase in enzyme concentration until substrate saturation is reached, hence it is assumed that increased flux through ASNN hydrolysis reaction implies higher L-asparaginase concentration, and strategies are discussed in accordance with this hypothesis. One of the predicted reaction amplification targets directly connected with the ASNN reaction is the reaction catalyzed by cytochrome C oxidase (represented as CYOO3 in Fig.2). The possible explanation for the prediction of CYOO3 reaction as a potential amplification target lies in water being generated as a by-product of the CYOO3 reaction. Water is essential for the hydrolytic reaction of ASNN and hence, an increase in CYOO3 flux increases the flux of water towards the ASNN reaction. This leads to an increase in L-asparaginase enzyme production due to the reason that an increase in water content leads to enhanced enzyme activity in hydrolytic reactions [34]. While the three reactions catalyzed by fructose biphosphate aldolase (FBA), glyceraldehyde 3 phosphate dehydrogenase (GAPD), and cytochrome c reductase (CYOR3m) are predicted as amplification targets as they are involved at the upstream of CYOO3 reaction ultimately increasing the flux through ASNN reaction. Prediction of reaction catalyzed by enolase (ENO) can be explained on the same line of thought as reaction catalyzed by enolase generates water as a byproduct. Apart from these reactions, the reaction catalyzed by glutamine synthetase (GLNS) catalyzing the conversion of amino acid glutamate and ammonia to glutamine is also identified as an amplification target. Amplifying this reaction reduces the accumulation of ammonia thereby driving the ASNN reaction forward increasing the flux through ASNN leading to an increase in L-asparaginase production. This result is in accordance with the role of glutamine synthetase whose expression increases as a result of ammonia deprivation, in the synthesis of L-asparaginase as observed in *Klebsiella aerogenes* [35].

#### 3.2.2 OptKnock to identify reaction knockout targets for ASNN production

OptKnock was carried out as mentioned in section 2.6 by setting different values of *Bacillus subtilis* growth rate. From the data obtained (tabulated in Table 3) it was seen that the highest flux through ASNN is achieved when a specific growth rate is set at the lowest value of 0.118 h^-1^. As this study aims at designing strategies for the overproduction of L-asparaginase, OptKnock results are analyzed further for the specific growth rate of 0.118 h^-1^. Due to the default output settings of the OptKnock function, five reactions are identified as knockout targets leading to an increase in ASNN flux. These are tabulated in Table 4 along with their complete name and the reaction formulae of each. OptKnock predicts a flux value of 109.33 mmol gDW^-1^ h^-1^ through ASNN reaction at the set specific growth rate and media conditions. Of the predicted five reactions, however, only two reactions catalyzed by cytidine deaminase (CYTD) and CMP deaminase (CMPDAi) are seen to be connected to the ASNN reaction as is clear from the network illustrated in Fig.4. Cytidine deaminase is involved in the pyrimidine salvage pathway that catalyzes the hydrolytic conversion of cytidine to uridine forming ammonia as a by-product while the enzyme CMP deaminase catalyzes the formation of uridine monophosphate (UMP) from cytidine monophosphate (CMP) with the release of ammonia. The prediction of these reactions as knockout targets can be argued on the terms that ammonia at higher concentrations proves toxic to the cell [36]. Thus, knocking out these reactions reduces ammonia accumulation in cells, eliminating its toxic effect on cell growth. Another probable reason could be that deaccumulation of ammonia due to deletion of these reactions, might drive the ASNN reaction in the forward direction. On analyzing the results of gene essentiality studies (details not mentioned); it was found that both CYTD and CMPDAi reactions are not essential for the growth of *Bacillus subtilis!*, hence knocking out these reactions does not affect the host cell growth rate.

**Fig.4.**
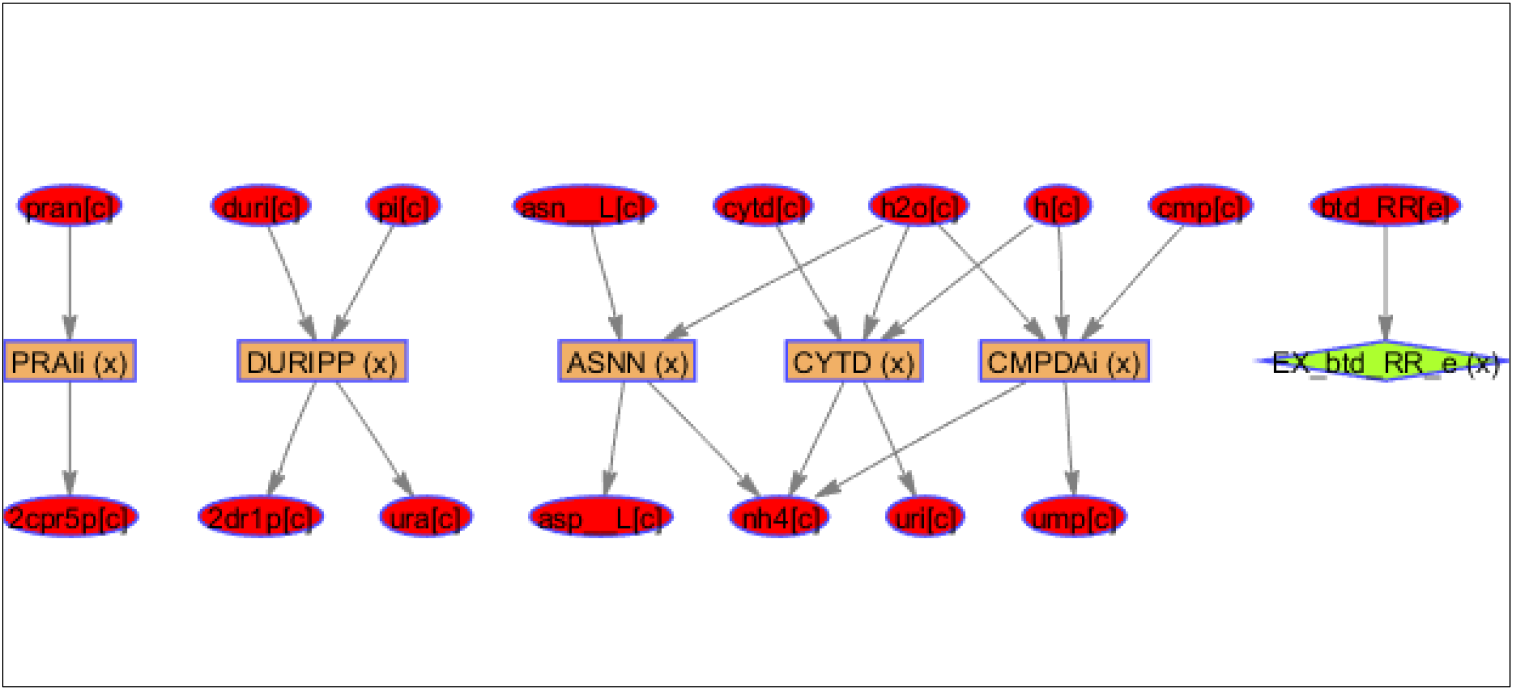
Biograph of metabolic network representing pathway connections of ASNN reaction with the identified knockout targets as generated by MATLAB R2019b in hierarchical layout

**Table 3:** The table lists flux through ASNN reaction (mmol g^-1^ h^-1^) for different specific growth rate values as predicted by OptKnock

| Specific growth rate set<br>by OptKnock method<br>(/hr) | Flux obtained through<br>ASNN reaction<br>(mmol/g/h) |
| --- | --- |
| 0.118 | 109.33 |
| 0.2 | 102.03 |
| 0.3 | 93.04 |
| 0.4 | 84.06 |
| 0.5 | 75.07 |
| 0.6 | 66.08 |

**Table 4:** Reaction knockout targets identified by OptKnock at the specific growth rate of 0.118 h^-1^ for maximizing flux through native L-asparaginase reaction. The table contains reaction ID as present in the model, complete reaction name, and reaction formulae

| Reaction ID | Reaction name | Reaction formula |
| --- | --- | --- |
| EX_btd_RR_e' | 'R R 2 3 Butanediol exchange' |  |
| 'CYTD' | 'Cytidine deaminase' | $\text{cytd}[\text{c}] + \text{h}_2\text{o}[\text{c}] + \text{h}[\text{c}] \rightarrow \text{nh}_4[\text{c}] + \text{uri}[\text{c}]$ |
| 'CMPDAi' | 'CMP deaminase' | $\text{cmp}[\text{c}] + \text{h}_2\text{o}[\text{c}] + \text{h}[\text{c}] \rightarrow \text{nh}_4[\text{c}] + \text{ump}[\text{c}]$ |
| 'DURIPP' | 'Deoxyuridine phosphorylase' | $\text{duri}[\text{c}] + \text{pi}[\text{c}] \rightleftharpoons 2\text{dr1p}[\text{c}] + \text{ura}[\text{c}]$ |
| PRAi' | 'Phosphoribosylanthranilate isomerase (irreversible)' | $\text{pran}[\text{c}] \rightleftharpoons 2\text{cpr5p}[\text{c}]$ |

While section (3.2) deals with designing strategies for increasing the production of native L-asparaginase in *Bacillus subtilis* host cell by identifying amplification and knockout targets, the following section details a study of overproduction of the heterologous L-asparaginase (rASNN) expressed in *Bacillus subtilis* host cell. The results are initially discussed for FBA, FVA, and robustness analysis after the addition of heterologous enzyme metabolic reaction to the GSMM as detailed in section 2.7 prior to discussing the strain improvement strategies.

### 3.3 *Bacillus subtilis* GSMM iYO844 analysis for heterologous L-asparaginase (rASNN) production

Integration of the heterogenous L-asparaginase gene in the form of a metabolic reaction (section 2.7) was attempted to simulate the production of recombinant L-asparaginase (rASNN) in the *Bacillus subtilis* host. A total of 3 reactions representing the production of rASNN and exchange reaction and 2 metabolites (intracellular and extracellular rASNN) were added to the model making the total reactions in the model to 1253 and total metabolites to 992.

#### 3.3.1 Flux balance analysis (FBA)

The addition of reactions to the model did not impair the growth of the organism. Even after reaction integration, the model predicted a specific growth rate of 0.6242 h^-1^ under given glucose and oxygen constraints. The growth rate decreased after constraining flux through the rASNN reaction.

#### 3.3.2 Flux variability analysis (FVA)

FVA predicts the minimum flux through the recombinant L-asparaginase (rASNN) reaction to be 0 mmol gDW^-1^ h^-1^ while the maximum flux allowed was 0.4028 mmol gDW^-1^ h^-1^.

#### 3.3.3 Robustness analysis

The growth rate drops linearly as flux through the rASNN reaction gradually increases as seen by the robustness analysis plot (Fig.5). This can be explained as both biomass growth and rASNN reactions compete for the same amino acids present in the cell. Robustness analysis helps in achieving a trade-off between the growth rate and recombinant L-asparaginase production rate. The data is further used for setting a specific growth rate value while identifying knockout strategies by the OptKnock method.

**Fig.5.**
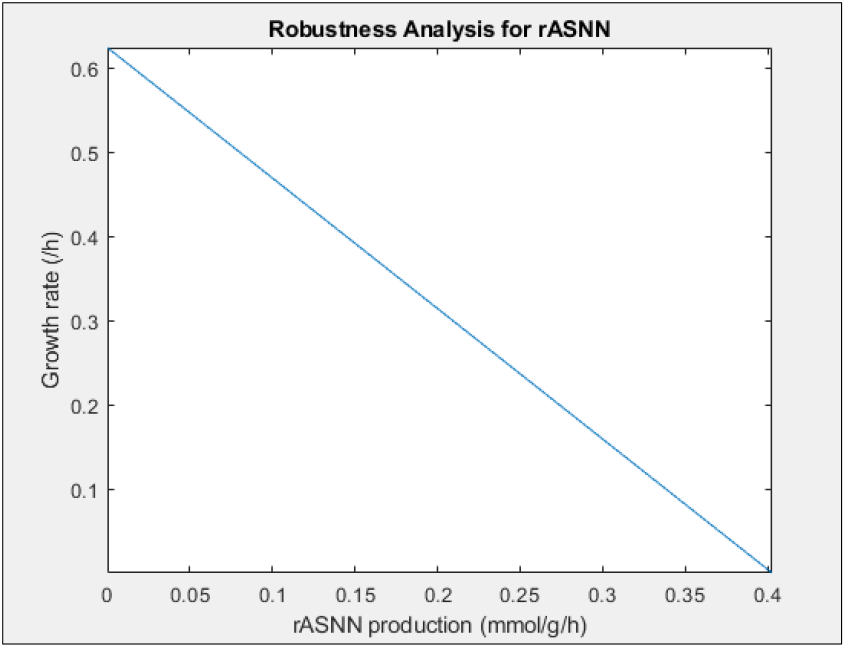
Plot representing the change of specific growth rate of *Bacillus subtilis* (h^-1^) with increase in rASNN production (mmol g^-1^ h^-1^) as obtained from robustness analysis. Growth rate declines linearly with increase in rASNN flux

### 3.4 Strain design for recombinant L-asparaginase (rASNN) production

#### 3.4.1 FSEOF to identify reaction amplification targets for rASNN production

FSEOF identifies reactions for which the flux amplifies when flux through rASNN is gradually increased. In accordance with the results obtained from robustness analysis, the specific growth rate of *Bacillus subtilis* is seen to decrease as flux through the rASNN reaction is gradually increased (Fig.6). The FSEOF method increased flux through the following listed reactions as flux through rASNN was increased from 0 to 0.4 mmol gDW^-1^ h^-1^ with an interval of 0.1. The reaction amplification targets predicted by FSEOF are summarized in the table below (Table 5) while Fig.7 represents the connection of these targets with that of rASNN in a metabolic network.

**Fig.6.**
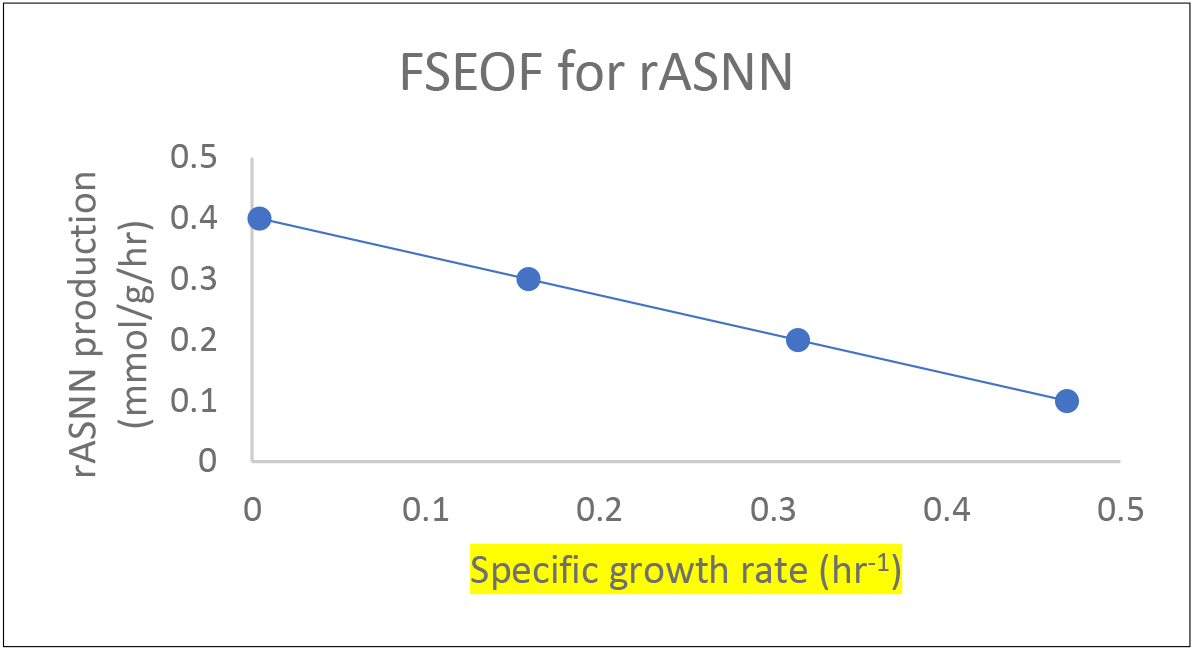
Plot representing the change in specific growth rate of *Bacillus subtilis* (h^-1^) as flux through rASNN reaction (mmol g^-1^ h^-1^) is gradually increased from the minimum to maximum limit as predicted by FVA. Growth rate increases linearly as rASNN flux drops, maximizing at 0 rASNN production

**Fig.7.**
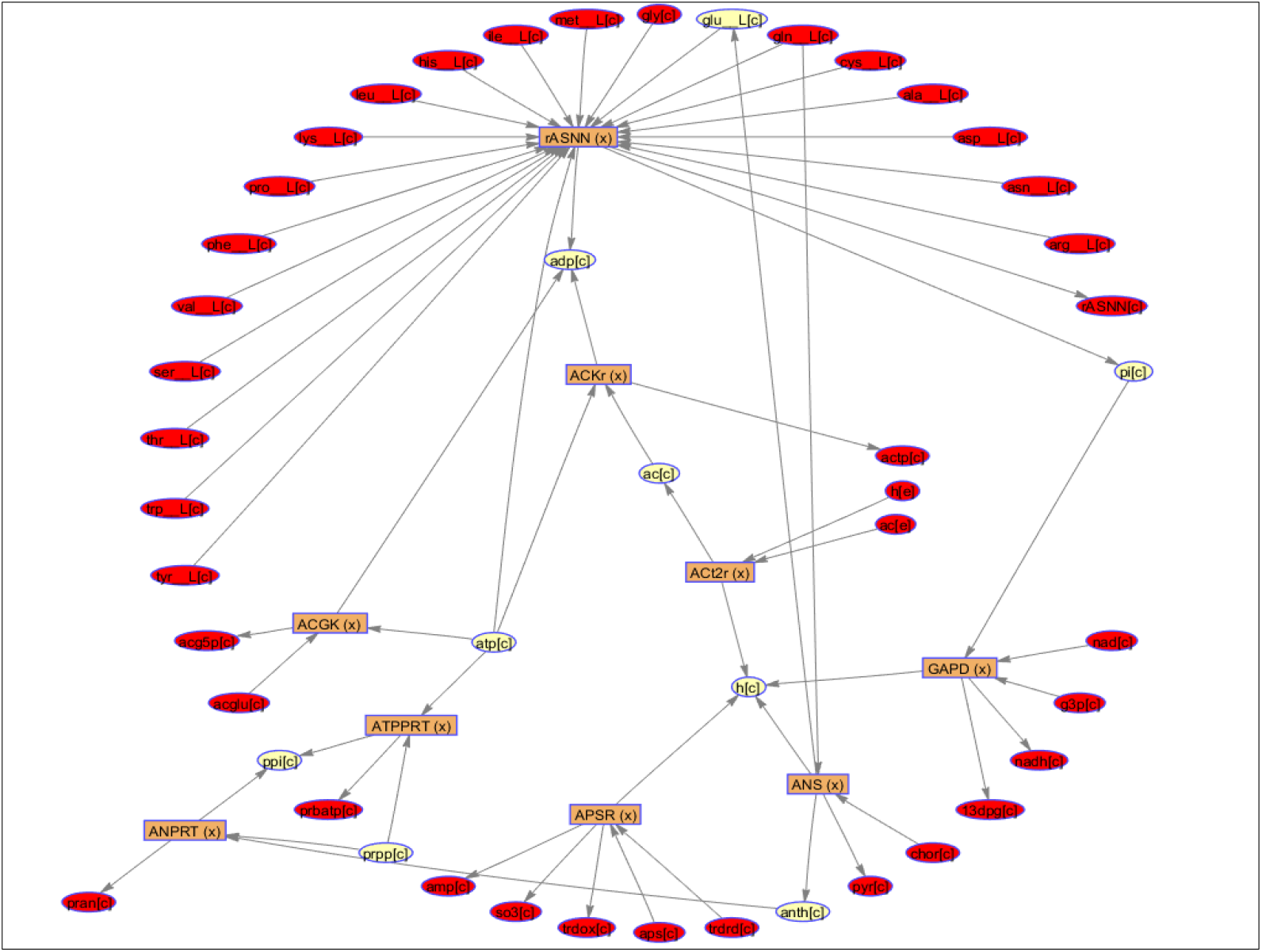
Biograph of metabolic network representing pathway connections of rASNN reaction with the identified amplification targets as generated by MATLAB R2019b in radial layout

**Table 5:**
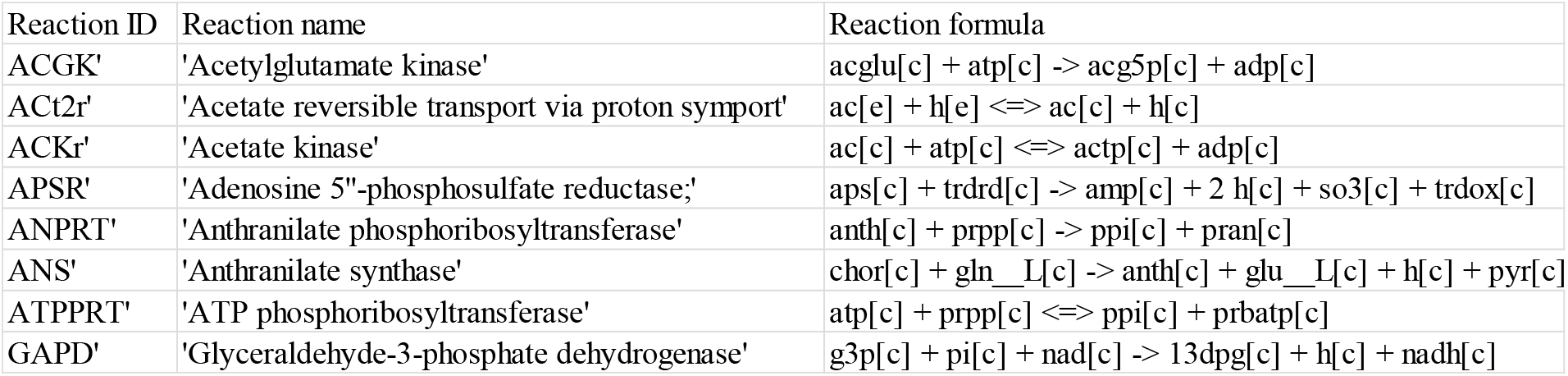
Reaction amplification targets identified by FSEOF for maximizing flux through recombinant L-asparaginase reaction. The table contains reaction ID as present in the model, complete reaction name, and reaction formulae

Out of all the possible amplification targets identified, the reactions catalyzed by glyceraldehyde-3-phosphate dehydrogenase (GAPD) and anthranilate synthase (ANS) are metabolically connected to rASNN production as is depicted in Fig.7. GAPD is involved in the glycolytic pathway and catalyzes the oxidative phosphorylation of glyceraldehyde-3-phosphate to 3-phospho-glycerol phosphate forming NADH as a byproduct in the process. Recombinant protein production in a host cell requires higher energy which is attempted by increased NADH production which is further used for ATP generation [37]. Amplification of GAPD reaction increases the flux towards NADH production in order to meet enhanced energy demand on cell as a result of metabolic stress from recombinant protein production. On the other hand, anthranilate synthase reaction belongs to the L-tryptophan biosynthesis pathway catalyzing the formation of anthranilate from L-glutamine and chorismate with pyruvate and L-glutamate as the by-products. Since L-glutamate is a required substrate for the formation of recombinant L-asparaginase, overexpression of ANS reaction increases the flux of L-glutamate towards rASNN reaction. Although amplification of ANS increases the flux of L-glutamate, it also consumes L-glutamine in the process which might lead to a decrease in L-glutamine flux towards rASNN. This problem can be overcome by providing L-glutamine externally in the media.

#### 3.4.2 OptKnock to identify reaction knockout targets for rASNN production

Similar to the case of *in-silico* production of native L-asparaginase, OptKnock was carried out as mentioned in section 2.7 by setting different values of *Bacillus subtilis* growth rate. From the data obtained (tabulated in Table 6) it was seen that the highest flux through ASNN is achieved when a specific growth rate is set at the lowest value of 0.118 h^-1^. As this study aims at designing strategies for the overproduction of L-asparaginase, OptKnock results are analyzed further for the specific growth rate of 0.118 h^-1^. Due to the default output settings of the OptKnock function, five reactions are identified as knockout targets leading to an increase in ASNN flux. These are tabulated in Table 7 along with their complete name and the reaction formulae of each. OptKnock predicts a flux value of 0.3266 mmol gDW^-1^ h^-1^ through rASNN reaction at the set specific growth rate and media conditions.

**Table 6:**
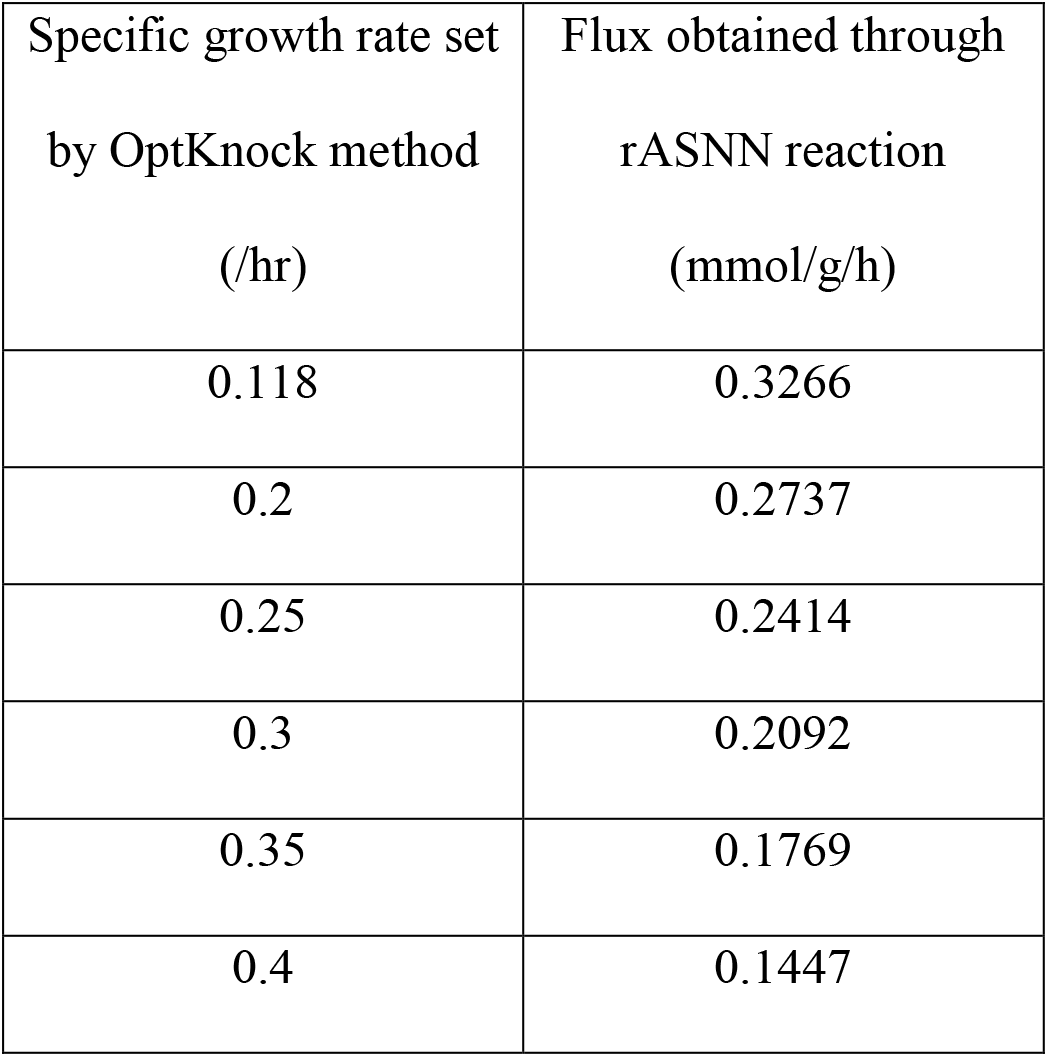
Table lists rASNN production rates (mmol g^-1^ h^-1^) for different specific growth rate values as predicted by OptKnock

| Specific growth rate set<br>by OptKnock method<br>(/hr) | Flux obtained through<br>rASNN reaction<br>(mmol/g/h) |
| --- | --- |
| 0.118 | 0.3266 |
| 0.2 | 0.2737 |
| 0.25 | 0.2414 |
| 0.3 | 0.2092 |
| 0.35 | 0.1769 |
| 0.4 | 0.1447 |

**Table 7:** Reaction knockout targets identified by OptKnock at the specific growth rate of 0.118 h^-1^ for maximizing the production of recombinant L-asparaginase reaction. The table contains reaction ID as present in the model, complete reaction name, and reaction formulae

| Reaction ID | Reaction name | Reaction formula |
| --- | --- | --- |
| 'ACGS' | 'N-acetylglutamate synthase' | accoa[c] + glu__L[c] -> acglu[c] + coa[c] + h[c] |
| 'CTPS1' | 'CTP synthase NH3 ' | atp[c] + nh4[c] + utp[c] -> adp[c] + ctp[c] + 2 h[c] + pi[c] |
| 'GART' | 'GAR transformylase-T' | atp[c] + for[c] + gar[c] -> adp[c] + fgam[c] + h[c] + pi[c] |
| 'DTMPK' | 'DTMP kinase' | atp[c] + dtmp[c] <=> adp[c] + dtdp[c] |
| 'NTD2' | '5''-nucleotidase (UMP)' | adp[c] + dtdp[c] -> pi[c] + uri[c] |

Out of the five targets identified, the reactions catalyzed by DTMP kinase (DTMPK, involved in both salvage and *de novo* pathways for deoxythymidine triphosphate biosynthesis), CTP synthase (CTPS1, involved in *de novo* CTP biosynthesis), and GAR transformylase (GART, involved in *de novo* purine biosynthesis) are considered potential targets as these reactions directly compete with rASNN reaction to utilize ATP (Fig.8). Knocking out these reactions will lead to an increase of ATP towards the rASNN reaction, increasing the rASNN production flux. Further, the reaction catalyzed by N-acetylglutamate synthase (ACGS, involved in arginine biosynthesis) requires L-glutamate as a substrate competing with the rASNN reaction for L-glutamate. Knocking out ACGS increases the flux of L-glutamate towards rASNN thereby increasing recombinant L-asparaginase production. According to gene essentiality analysis (details not mentioned), CTPS1, GART, and ACGS are found to be non-essential for biomass growth while DTMPK is considered to be an essential reaction for *Bacillus subtilis* growth due to its involvement in dTTP biosynthesis.

**Fig.8.**
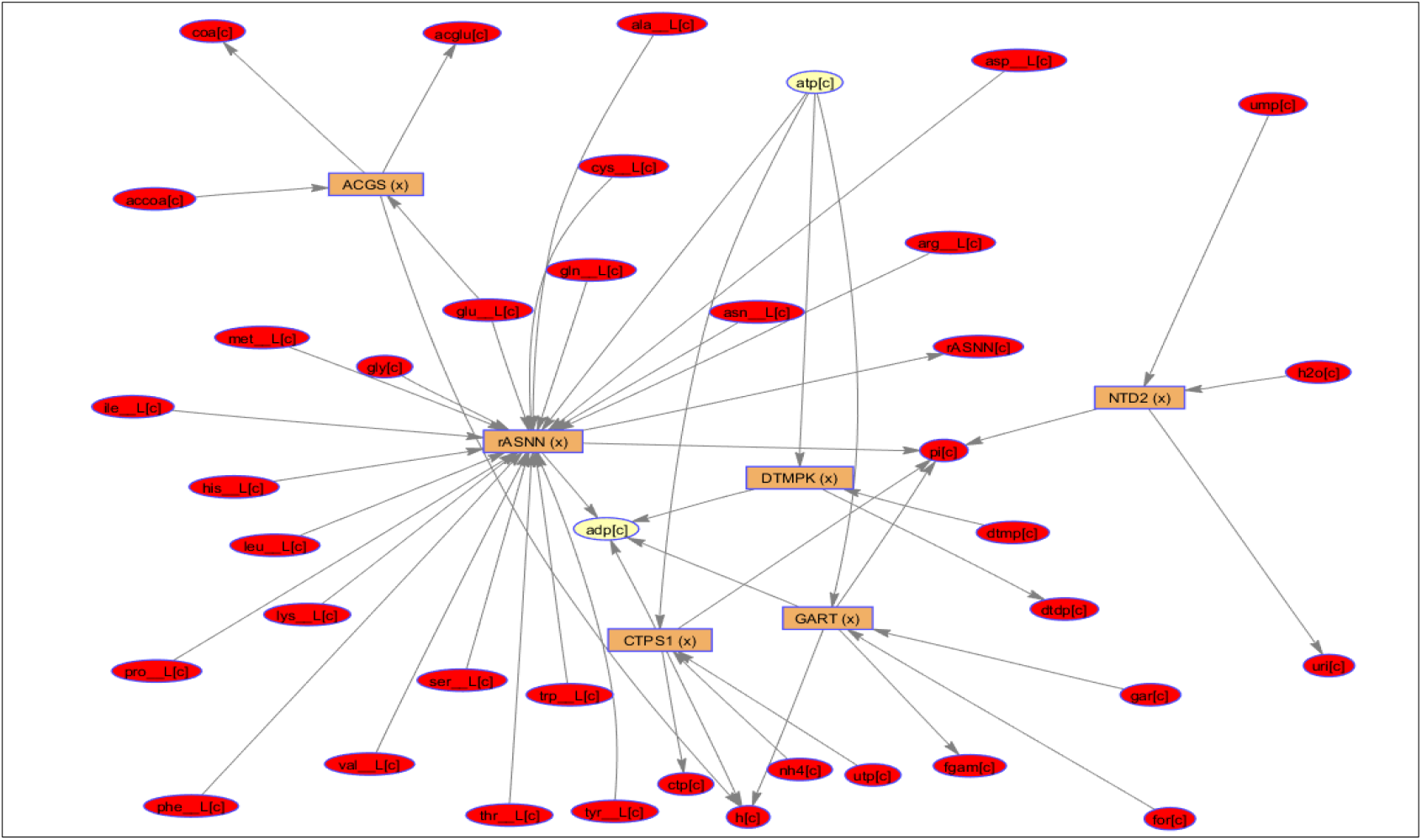
Biograph of metabolic network representing pathway connections of rASNN reaction with the identified knockout targets as generated by MATLAB R2019b in equilibrium layout

### 3.5 Strain design for recombinant L-asparaginase production with amino acid supplements in media

On providing the media with all 20 amino acids as mentioned in section 2.8, the model was utilized for the simulation of host cell growth and production of recombinant L-asparaginase. Due to the additional media components, both growth rate and rASNN flux were found to be increased. The maximum specific growth rate of 0.86 h^-1^ and maximum recombinant L-asparaginase production rate of 1.4 mmol gDW^-1^ h^-1^ was obtained after amino acids were externally supplied. It is clear from the plot of rASNN production rate and specific growth rate that as flux through rASNN increases, the growth rate starts to decline. The specific growth rate decreased gradually from 0.86 h^-1^ to 0.14 h^-1^ (Fig.9). Although the addition of amino acids in media leads to an increase in both growth rate as well as recombinant L-asparaginase production rate yet the strategy has to be economical as the addition of amino acids in media might lead to an increase in production cost. Further experimentation is required to arrive at a trade-off between higher production rates and economic feasibility.

**Fig.9.**
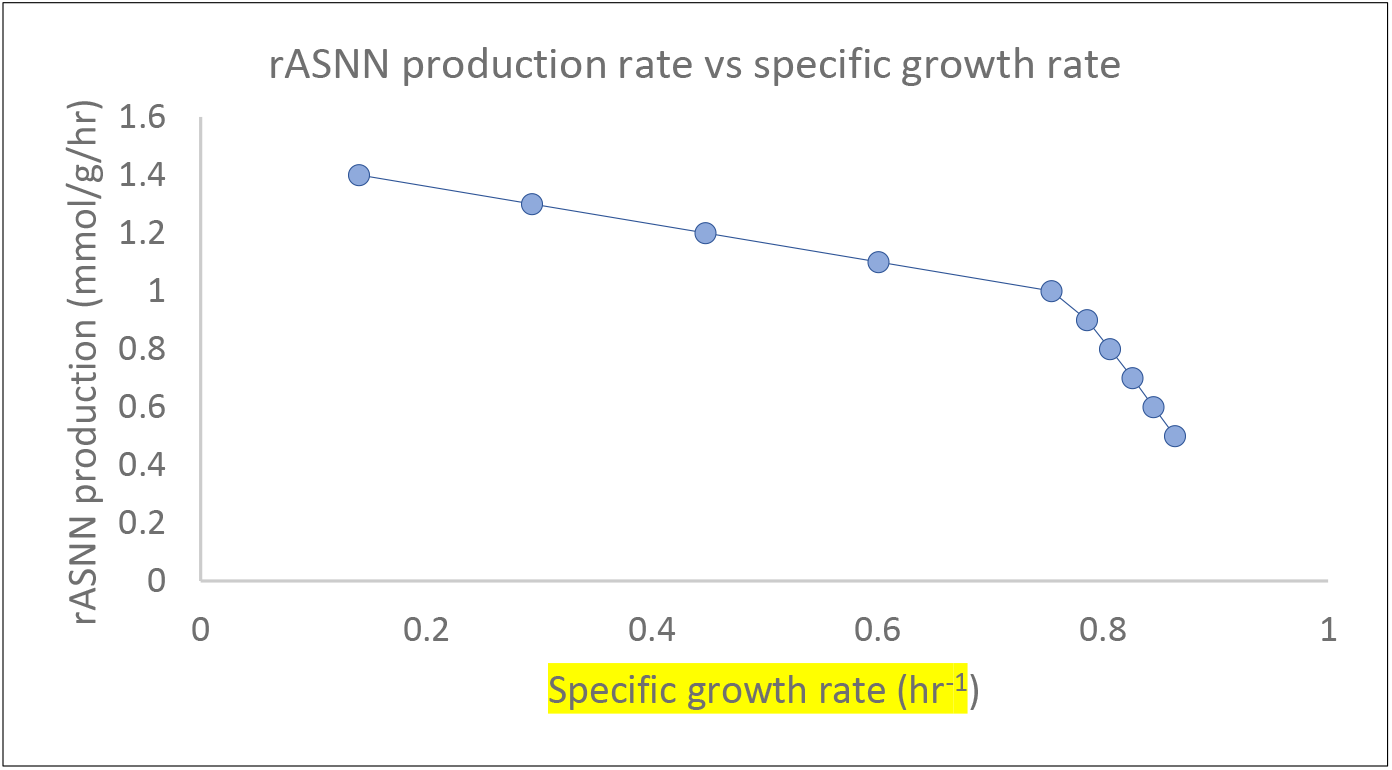
Plot representing the change of specific growth rate of Bacillus subtilis as rASNN production rate increases gradually after supplementing the media with amino acids. Growth declines almost linearly until 0.75 h^-1^ following a steep slope

## 4. Conclusion

GSMM, an important tool based on the principles of systems biology finds extensive applications in metabolic engineering and strain design for the production of industrially important products. This article details the analysis of GSMM of *Bacillus subtilis* for the production of L-asparaginase, a chemotherapeutic drug used in the treatment of ALL cancer. Analysis of GSMM gives a brief understanding of the metabolism of *Bacillus subtilis* which is further utilized to study the production of native L-asparaginase. Computational design methods like FSEOF and OptKnock are utilized to predict strategies to design model *Bacillus subtilis* producing higher L-asparaginase yields. Further, due to the limitations of commercially available L-asparaginase drug attributed to its glutaminase activity, a heterologous gene encoding a putative glutaminase-free L-asparaginase is added to GSMM of *Bacillus subtilis* in an attempt to simulate recombinant L-asparaginase production. Amplification and knockout strategies are then designed for higher production of recombinant L-asparaginase in the *Bacillus subtilis* host. The results obtained describe *in silico* strategies and further confirmation of the same by experimental studies is required to validate the results. It is further observed that providing amino acids as additional media components enhances recombinant L-asparaginase production as well as growth rate. This study reduces the efforts required for strain improvement by minimizing the targets to be identified and provides potential prerequisite information for further studies. Experimental validation of the obtained results remains to be performed. The study can help in economizing the process of effective L-asparaginase drug production.

## ABBREVIATIONS

ALL: Acute Lymphoblastic Leukemia
ASNN: L-asparaginase
COBRA: COnstraints Based Reconstruction and Analysis
FBA: Flux balance analysis
FSEOF: Flux Scanning based on Enforced Objective Flux
FVA: Flux variability analysis
GSMM: Genome-scale metabolic model
NAD: Nicotinamide adenine dinucleotide
NADH: Nicotinamide adenine dinucleotide-hydrogen (reduced form of NAD)
PEG: Polyethylene glycol
rASNN: recombinant L-asparaginase
SBML: Systems Biology Markup Language

## 5. Author Statement

N. S. Barge: Conceptualization, Methodology, Validation, Investigation, Data Curation, Writing - Original Draft. A. Sahoo: Formal Analysis, Writing - review & editing. V.V. Dasu: Conceptualization, Resources, Supervision, Visualization, Writing - review & editing.

## 7. Acknowledgement

The authors are grateful to IIT Guwahati and the Ministry of Education, Govt. of India for the financial support to carry out this work.

