## Supplementary file S1 for "Analysis of the genome-scale metabolic model of *Bacillus subtilis* to design novel in-silico strategies for native and recombinant L-asparaginase overproduction"

**1. Methodology**

**1.1** **Genome scale metabolic model analysis:**

The COBRA Toolbox is initiated using the following function:

>> initCobraToolbox

The model is loaded by the following function and choosing the required file from directory:

>> model = readCbModel;

Following functions were used to change the bounds for glucose and oxygen. Negative sign for values here, signifies uptake of the media constituents. The reactions are identified by the syntax present in the model.

>> model = changeRxnBounds (model, ‘EX_glc_D_e’, -15, ‘l’);

>> model = changeRxnBounds (model, ‘EX_o2_e’, -20, ‘l’);

**1.2 Flux balance analysis (FBA):**

FBA is performed using ‘optimizeCbModel’ function:

>> FBAsolution = optimizeCbModel (model, ‘max’);

The function is used to maximize or minimize the objective function using ‘max’ or ‘min’ respectively. COBRA Toolbox also allows listing of the non-zero flux values along with their reaction IDs for easier analysis of flux distribution.

>> fluxData = FBAsolution.v;

>> nonZeroFlag = 1;

>> printFluxVector (model, fluxData, nonZeroFlag)

**1.3 Flux variability analysis (FVA):**

To perform simultaneous FVA for all the reactions in the model, following command is used:

>> [minFlux, maxFlux] = fluxVariability (model);

>> FVA = table (model.rxns, minFlux, maxFlux)

The last command generates a table listing all the reactions in the model represented by the unique reaction IDs and the respective minimum and maximum flux a particular reaction achieves under the specific media conditions. FVA for native L-asparaginase (ASNN) reaction to determine the range of fluxes the reaction can achieve is performed as detailed below:

>> FBAsolution = optimizeCbModel (model, ‘max’);

>> model = changeRxnBounds (model, ‘BIOMASS_BS_10’, FBAsolution.f, ‘b’);

>> model = changeObjective (model, ‘ASNN’);

>> FBAsolutionMax = optimizeCbModel (model, ‘max’);

>> FBAsolutionMin = optimizeCbModel (model, ‘min’);

**1.4 Robustness analysis:**

Prior to performing robustness analysis, native L-asparaginase reaction is set as the control reaction as we wanted to assess the effect of flux change in L-asparaginase reaction on that of growth rate of *Bacillus subtilis*.

>> controlRxn = ‘ASNN’;

>> [controlFlux, objFlux] = robustnessAnalysis (model, controlRxn);

**1.5** **Identification of gene amplification targets using flux scanning based on enforced objective flux (FSEOF) method:**

The flux through ASNN reaction is gradually increased from 0 to 37 mmol gDW^-1^ h^-1^ with an interval of 5. The flux distribution for all ASNN values is scanned to determine the reactions which are gradually increasing and considered as reaction amplification targets.

>> model = changeRxnBounds (model, ‘ASNN’, 5, ‘l’);

>> model = changeRxnBounds (model, ‘ASNN’, 10, ‘l’);

>> model = changeRxnBounds (model, ‘ASNN’, 15, ‘l’);

>> model = changeRxnBounds (model, ‘ASNN’, 20, ‘l’);

>> model = changeRxnBounds (model, ‘ASNN’, 25, ‘l’);

>> model = changeRxnBounds (model, ‘ASNN’, 30, ‘l’);

>> model = changeRxnBounds (model, ‘ASNN’, 35, ‘l’);

>> model = changeRxnBounds (model, ‘ASNN’, 37, ‘l’);

**1.6** **Identification of gene knockout targets using OptKnock:**

The model is loaded into MATLAB by initiating COBRA Toolbox and specifying glucose and oxygen constraints as discussed earlier (section 1.1).

>> options.targetRxn = ‘ASNN’; (sets the target reaction to increase flux through native L-asparaginase)

>> constrOpt.rxnList = {‘BIOMASS_BS_10}; (sets biomass reaction to fix value)

>> constrOpt.values = [0.118]; (sets biomass growth value)

[optKnockSol, bilevelMILPproblem] = OptKnock (model, model.rxns, options, constrOpt);

The interconnection of the target reactions with that of ASNN is visualized using in-built visualization functions.

**1.7 Integration of heterologous L-asparaginase production metabolic reaction into iYO844:**

The gene for L-asparaginase was added to *Bacillus subtilis* iYO844 in the form of a reaction based on the amino acid sequence of *Rhizomucor miehei* L-asparaginase. The amino acid sequence of *Rhizomucor miehei* L-asparaginase was obtained from UniProt (https://www.uniprot.org/uniprot/W0G253) and is as follows:

MDSRTTAHVPIYDNAAVEHQLRADRDEMLAPDFSRVLVIYTGGTIGMKHTPEHGYIPLPNYLAQSLARLIRFHDPSHGFLSRSSSQENHADGVTLKEDFTRITNMVRQVQPSGETVLAHLPSLITPVSLYGKRIRYSILEYDPLLDSCNITMDDWVRIARDIEANYEYFDAFIVLHGTDTMAYTASALSFMLEELGKTVIITGSQVPLTEVRNDAVENLLGALTIAGHFVIPEVCLYFGDKLYRGNRTSKISAVDFDAFDSPNLPALVNLGIDIDVKWPLVLRPTHIAKFRSHKVLNRNVASLRLFPGINESTVRAFLAPPLQGVVLETYGAGNAPARQGLLAALKEACDRGVVIVNCTQCRKGLVTDSYATGRQLAAIGVVAGADMTPECALTKLSYLLGKIPDDPQKVRLLMTRNLRGELTVRAEKQRFSASGSRSQLLLDIFANISARGKVAAKTHVDQQIMSVEEEQLAEKMLAPMLLCSAASANDVQSMKMLSDAMGDILNLNCVDYAGRSPLHIACRDGHIAIVEYLLLHGASVHVRDRWGHTPLFVAVVGKHAQVVSMLRRAGAHLSVNEQSDMGPAWLKAVRDNDVEFVKIALEAGWPVNWAEPVEGRRAIDIAVCYGRVDLLKLLLLQVDCKTDQPDRWGFTIIDKLSLLEKQEKKSIDPETLKEIAQLLGKE

The table below specifies the % content of each amino acid in *Rhizomucor miehei* L-asparaginase based on its amino acid sequence and the coefficient of each amino acid used to design the metabolic reaction for recombinant enzyme to be integrated into *Bacillus subtilis* GSMM.

Table S1:

| Amino acid | % protein (w/w) | MW (g/mol) | mmol/g protein |
| --- | --- | --- | --- |
| Alanine | 6.136 | 71.09 (65) | 0.8631 |
| Arginine | 8.918 | 156.20 (43) | 0.5709 |
| Asparagine | 3.637 | 114.12 (24) | 0.3187 |
| Aspartate | 6.877 | 115.10 (45) | 0.5975 |
| Cysteine | 1.507 | 103.16 (11) | 0.1461 |
| Glutamate | 5.956 | 128.15 (35) | 0.4648 |
| Glutamine | 3.944 | 129.13 (23) | 0.3054 |
| Glycine | 3.258 | 57.07 (43) | 0.5709 |
| Histidine | 3.642 | 137.16 (20) | 0.2655 |
| Isoleucine | 5.861 | 113.18 (39) | 0.5178 |
| Leucine | 12.173 | 113.18 (81) | 1.0755 |
| Lysine | 5.106 | 128.19 (30) | 0.3983 |
| Methionine | 2.962 | 131.21 (17) | 0.2257 |
| Phenylalanine | 3.713 | 147.19 (19) | 0.2522 |
| Proline | 3.998 | 97.13 (31) | 0.4116 |
| Serine | 4.741 | 87.09 (41) | 0.5444 |
| Threonine | 4.565 | 101.12 (34) | 0.4514 |
| Tryptophan | 1.731 | 186.23 (7) | 0.0929 |
| Tyrosine | 3.900 | 163.19 (18) | 0.2390 |
| Valine | 7.373 | 99.15 (56) | 0.7436 |

The reactions were added using ‘addReaction’ in-built function available in COBRA Toolbox.

>> model = addReaction(model,'rASNN','metaboliteList',{'ala__L[c]','arg__L[c]','asn__L[c]','asp__L[c]','cys__L[c]','glu__L[c]','gln__L[c]','gly[c]','his__L[c]','ile__L[c]','leu__L[c]','lys__L[c]','met__L[c]','phe__L[c]','pro__L[c]','ser__L[c]','thr__L[c]','trp__L[c]','tyr__L[c]','val__L[c]','atp[c]','rASNN[c]','adp[c]','pi[c]'},'stoichCoeffList',[-0.8631 -0.5709 -0.3187 -0.5975 -0.1461 -0.4648 -0.3054 -0.5709 -0.2655 -0.5178 -1.0755 -0.3983 -0.2257 -0.2522 -0.4116 -0.5444 -0.4514 -0.0929 -0.2390 -0.7436 -3.984 1 3.984 3.984], 'reversible',false);

The following reactions simulate the exchange and extracellular production of recombinant L-asparaginase:

>> model = addReaction(model,'rASNN[e]','metaboliteList',{'rASNN[c]','rASNN[e]'},'stoichCoeffList',[-1 1],'reversible',false);

>> model = addExchangeRxn(model, {'rASNN[e]'});

**1.8 Model analysis after integration of heterologous L-asparaginase production metabolic reaction into iYO844:**

*Flux variability analysis*: FVA for recombinant L-asparaginase (rASNN) reaction to determine the range of fluxes the reaction can achieve is performed as detailed below,

>> FBAsolution = optimizeCbModel (model, ‘max’);

>> model = changeRxnBounds (model, ‘BIOMASS_BS_10’, FBAsolution.f, ‘b’);

>> model = changeObjective (model, ‘rASNN’);

>> FBAsolutionMax = optimizeCbModel (model, ‘max’);

>> FBAsolutionMin = optimizeCbModel (model, ‘min’);

*Robustness analysis*: Prior to performing robustness analysis, recombinant L-asparaginase production reaction is set as the control reaction.

>> controlRxn = ‘rASNN’;

>> [controlFlux, objFlux] = robustnessAnalysis (model, controlRxn);

**1.9 FSEOF and OptKnock methods for overproduction of recombinant L-asparaginase enzyme:**

*FSEOF*: To identify the amplification targets, FSEOF is carried out by increasing the flux through rASNN reaction from 0 to 0.4028 mmol gDW^-1^ h^-1^ with an interval of 0.1. The flux distribution for all rASNN values is scanned to determine the reactions which are gradually increasing and considered as reaction amplification targets.

>> model = changeRxnBounds (model, ‘rASNN’, 0.1, ‘l’);

>> model = changeRxnBounds (model, ‘rASNN’, 0.2, ‘l’);

>> model = changeRxnBounds (model, ‘rASNN’, 0.3, ‘l’);

>> model = changeRxnBounds (model, ‘rASNN’, 0.4, ‘l’);

*OptKnock*: Knockout targets are identified by performing OptKnock and setting the biomass growth rate value to 0.118 h^-1^.

>> options.targetRxn = ‘rASNN’; (sets the target reaction to overproduce recombinant L-asparaginase)

>> constrOpt.rxnList = {‘BIOMASS_BS_10}; (sets biomass reaction to fix value)

>> constrOpt.values = [0.118]; (sets biomass growth value)

[optKnockSol, bilevelMILPproblem] = OptKnock (model, model.rxns, options, constrOpt);

The interconnection of the target reactions with that of rASNN is then visualized using in-built visualization functions.

**1.10 Adding amino acids to the media for increasing rASNN production:**

The uptake rates are set as follows:

>> model = changeRxnBounds(model,'EX_ala__L_e',-0.1,'l');

>> model = changeRxnBounds(model,'EX_arg__L_e',-0.55,'l');

>> model = changeRxnBounds(model,'EX_asn__L_e',-0.1,'l');

>> model = changeRxnBounds(model,'EX_asp__L_e',-2,'l');

>> model = changeRxnBounds(model,'EX_cys__L_e',-0.39,'l');

>> model = changeRxnBounds(model,'EX_glu__L_e',-0.1,'l');

>> model = changeRxnBounds(model,'EX_gln__L_e',-2,'l');

>> model = changeRxnBounds(model,'EX_gly_e',-0.3,'l');

>> model = changeRxnBounds(model,'EX_his__L_e',-0.1,'l');

>> model = changeRxnBounds(model,'EX_ile__L_e',-0.3,'l');

>> model = changeRxnBounds(model,'EX_leu__L_e',-0.45,'l');

>> model = changeRxnBounds(model,'EX_lys__L_e',-0.3,'l');

>> model = changeRxnBounds(model,'EX_met__L_e',-0.16,'l');

>> model = changeRxnBounds(model,'EX_phe__L_e',-0.17,'l');

>> model = changeRxnBounds(model,'EX_pro__L_e',-0.2,'l');

>> model = changeRxnBounds(model,'EX_ser__L_e',-0.5,'l');

>> model = changeRxnBounds(model,'EX_thr__L_e',-0.9,'l');

>> model = changeRxnBounds(model,'EX_trp__L_e',-0.05,'l');

>> model = changeRxnBounds(model,'EX_tyr__L_e',-0.14,'l');

>> model = changeRxnBounds(model,'EX_val__L_e',-0.4,'l');

**1.11 Visualization of metabolic networks:**

Initially a cell vector containing the reaction IDs of reactions to be included in the layout is specified.

>> rxns = {cell array of reactions to be visualized};

The following function is used for pathway visualization,

>> [Involved_mets, Dead_ends] = draw_by_rxn (model, rxns, ‘true’)
